# Mammalian orthoreovirus infection in human epidermal growth factor receptor 2 positive (HER2+) breast cancer cells

**DOI:** 10.1101/2023.05.10.540250

**Authors:** Nicole A. Jandick, Nicolette Kirner, Cathy L. Miller

## Abstract

Mammalian orthoreovirus (MRV) is a clinically benign oncolytic virus which has been investigated for use in multiple cancer types, including breast cancer (BC). In human clinical trials, MRV has been shown to be safe, and multiple BC patients have shown partial responses to intratumoral and intravenous virus delivery. Combination therapies inclusive of MRV and current FDA approved BC chemotherapies are being investigated to target metastatic, early BC, and triple negative BC. Though MRV is being tested clinically, we still do not fully understand the highly variable patient responses to MRV therapy. One of the most aggressive BC subtypes is HER2+ BC, in which human epidermal growth factor receptor 2 (HER2) is dysregulated, resulting in increased growth, survival, and metastasis of cancer cells. FDA approved therapies, trastuzumab and pertuzumab, target HER2 to prevent signaling of the phosphoinositide 3-kinase (PI3K) pathway. However, recent findings show that accumulation of hypoxia inducible factor-1 alpha (HIF-1α) in HER2+ BC cells contributes to trastuzumab resistance. In this work, we provide evidence that MRV infects, replicates in, and kills HER2 overexpressing cells. MRV infection is also found to have variable effects on signaling pathways that activate or are activated by HER2 expression. Finally, we show that MRV reduces HIF-1α accumulation in all the cell lines tested, including a HER2+ BC cell line. These studies provide further evidence that MRV holds promise for use in conjunction with trastuzumab to treat HER2+ BC patients.

## Introduction

Breast cancer (BC) is the most commonly diagnosed cancer globally with incidence rates steadily climbing since the mid-2000s [1]. Overall BC has a 90% 5-year survival rate, however, this rate varies greatly with BC stage and molecular subtype of the tumor [1]. There are four main subtypes of BC, luminal A, luminal B, human epidermal growth factor receptor positive (HER2+) and triple negative (TNBC). Luminal A tumors are positive for estrogen receptor (ER) and either positive or negative for progesterone receptor (PR), exhibit no HER2 expression and have low expression of the nuclear proliferation marker, Ki-67 [2]. Luminal B tumors are ER positive, PR positive or negative, HER2 negative and exhibit high Ki-67 expression [2]. Tumors classified as HER2+, overexpress HER2 and are branched into two subgroups, luminal HER2, which include tumors that are ER+ and PR+ and express 15-30% Ki-67, and HER2-enriched, which are ER and PR negative, and express over 30% Ki-67 [2]. The fourth subtype is triple negative BC (TNBC), which is ER, PR and HER2 negative. TNBC has the lowest survival rate, due to lack of response to hormone therapy and HER2+ BC has the second lowest survival rate [3].

Overexpression of HER2 is seen in 15-30% of BC patients [4]. HER2 overexpression occurs because of increased copies and/or increased transcription of the HER2 gene, both of which result in increased HER2 receptors on the cell surface [4]. Members of the HER family signal through the formation of homo- or heterodimers between HER1-HER4. The preferential dimerization of HER2 and HER3 signals through activation of the phosphoinositide 3-kinase (PI3K) pathway, leading to increased cell survival and tumor proliferation [5]. This pathway leads to increased protein translation via activation of 3-phosphoinositide kinase-1 (PDK1), which phosphorylates the Thr(308) site on Akt [6]. Additional Akt phosphorylation at Ser(473) by other kinases such as mammalian target of rapamycin complex 2 (mTORC2) increases mTOR1 activity, resulting in inhibition of eukaryotic translation initiation factor 4E binding protein 1 (4E-BP1) by phosphorylation and activation of 40S ribosomal S6 kinase (S6K) by dephosphorylation, increasing protein translation. Currently two immunotherapies, trastuzumab and pertuzumab are approved for use in HER2+ BC patients. Both therapies are humanized antibodies that work to bind HER2 and inhibit dimerization and downstream signaling pathways, including PI3K [7].

Trastuzumab in combination with taxanes is the standard of care treatment for HER2+ BC patients. With the discovery of trastuzumab, HER2+ BC survival rate increased significantly [8]. Trastuzumab is thought to function by binding subdomain IV of the HER2 extracellular domain, preventing homodimerization with other HER2 proteins, preventing downstream signaling pathways, and ultimately leading to decreased growth of HER2+ cells [9]. Additionally, trastuzumab is known to induce antibody dependent cell cytotoxicity (ADCC) [10, 11] and reduce HER2 shedding by inhibiting metalloproteinase activity [12]. Although trastuzumab inhibits additional HER2 shedding that generates a truncated form of HER2, p95HER2, cells that already express p95HER2 cannot be recognized by trastuzumab. Additionally, p95HER2 can form from internal translation initiation sites on HER2 at amino acid (aa) 611 or 678 [13]. Unfortunately, expression of p95HER2 is associated with resistance to trastuzumab therapy [14]. Moreover, recent studies have found that accumulation of hypoxia inducible factor-1 alpha (HIF-1α) can also lead to trastuzumab resistance through a pathway that leads to downregulation of phosphatase and tensin homolog (PTEN) [15]. Pertuzumab, like trastuzumab, is a humanized antibody that can recognize HER2, however it binds subdomain II of the extracellular domain of HER2. This prevents heterodimerization of HER2 and blocks ligand dependent signaling pathways [16]. Pertuzumab has also been shown to induce ADCC [17, 18]. Currently pertuzumab is given in combination with trastuzumab treatment in patients with HER2+ BC. The development of resistance to trastuzumab and pertuzumab, makes the development of new therapies to treat HER2+ BC patients imperative.

Mammalian orthoreovirus (MRV) is a clinically benign oncolytic virus that has been studied preclinically and tested in over 30 clinical trials for the treatment of various cancer types [19]. While the effect of MRV on tumor cells has historically been considered primarily one of cell lysis and death, recent studies have highlighted the ability of the virus to induce the maturation of dendritic cells (DC) and increase the presentation of both viral and tumor associated antigens [20, 21]. This allows previously immune-suppressed, or cold tumor microenvironments to become ‘hot’ with active immune cells and stimulate an anti-tumor immune state [21]. MRV as a monotherapy has shown variable success in patients, with most patients exhibiting partial or no response and only rare patients exhibiting complete tumor regression. Our limited understanding of the MRV mechanism of action and of which tumor genetic types/subtypes and microenvironments are most susceptible to MRV, combined with the impracticality of an approach focused specifically on individual tumor types/subtypes in early clinical trials are likely contributing factors to the published variations in clinical responses to MRV therapy. Early pre-clinical studies found that MRV has enhanced infectivity of cells with an activated RAS pathway [22]. Further research showed that in specific cells (NIH 3T3 cells and H-RAS), an active RAS/RalGEF/p38 pathway is needed for susceptibility to MRV infection and oncolysis [23]. Activation of the epidermal growth factor receptor (EGFR/HER1) pathway, which can lead to activated RAS and signaling of the RAS/RalGEF/p38 pathway, has also been reported to aid in susceptibility of two mouse cell lines, NR6 and B82, to MRV infection [24, 25]. Additional studies found MRV does not bind EGFR/HER1 to enter cells, but instead activated EGFR/HER1 in cells enhances infection by a poorly understood mechanism [26]. EGFR/HER1 and HER2 are members of the epidermal growth factor receptor family, which can activate the same pathways depending on their dimerization partner, suggesting MRV infection in cells overexpressing HER2 may be enhanced.

Recently MRV has been given fast track designation for use in metastatic BC in combination with paclitaxel. This was granted in response to a clinical trial in which overall survival doubled in patients with TP53 mutations that were treated with MRV and paclitaxel compared to patients treated with paclitaxel alone [27]. However, when common benign polymorphisms of TP53 were excluded from the TP53 mutation population, the results inverted, with patients who received only paclitaxel showing increased overall survival compared to those treated with paclitaxel and MRV [27]. Interestingly, in this same trial, patients with Phosphatidylinositol-4,5-Bisphosphate 3-Kinase Catalytic Subunit Alpha (PI3KCA) wildtype (WT), Adenomatous Polyposis Coli (APC) WT, Kinase Insert Domain Receptor (KDR) mutations, or Mesenchymal Epithelial Transition (MET) mutations showed increased overall survival with MRV and paclitaxel treatment compared to paclitaxel alone [27]. This further supports that the genetic makeup of tumors likely plays a significant role in susceptibility to the therapeutic effects of MRV and underscores the idea that there is still much to learn about which tumor environments are most suited for MRV treatment.

In addition to genetic changes within tumors that may alter the therapeutic utility of MRV, the virus also modifies signaling pathways that are known to play significant roles in tumor growth and progression. One such pathway that is modified during MRV infection is the response to hypoxia, primarily regulated by the transcription factor HIF-1α. Under normal oxygen conditions HIF-1α is targeted for degradation by the proteasome in an oxygen-dependent manner. However, in hypoxic microenvironments which are routinely found in solid tumors, this pathway is not active and HIF-1α accumulates, leading to activation of signal transduction pathways that contribute to tumor cell proliferation and metastasis [28]. In hypoxic conditions, MRV infection has been shown to induce a massive decrease in the accumulation of HIF-1α. This phenotype, while seen in a large variety of cell types such as colon cancer cells [29], lung carcinoma cell lines [30, 31], prostate cancer cells [32-34], and a sarcoma xenograph in vivo model [35], is cell-type dependent as HIF-1α stabilization was seen in response to MRV infection of human glioblastoma cells [36]. The exact mechanism of MRV reduction in HIF-1α accumulation is not known. However, MRV inhibits the accumulation of HIF-1α independently of the oxygen-dependent pathway in a manner dependent on RACK1 and ubiquitin-mediated proteasomal degradation [32, 37]. Moreover, a step between viral capsid cleavage and transcription of viral mRNA during the MRV life cycle has been found to be sufficient to induce the MRV effect on HIF-1α accumulation [34]. Approximately 25-40% of BC tumors show accumulation of HIF-1α, which can lead to drug resistance and increased chance of recurrence [38-43]. Additionally, overexpression of HER2 has been shown to increase the accumulation of HIF-1α in normoxic conditions [44, 45]. In a mouse model where the murine HER2 homolog, Neu, is overexpressed (MMRV-Neu), HIF-1α has been found to be required for HER2-mediated tumor growth [46]. Finally, HIF-1α has been shown to play a role in development of trastuzumab resistance [15]. These findings suggest that there are links between HIF-1α and HER2 signaling regulation and may suggest that MRV modification of HIF-1α accumulation could influence HER2+ signaling in BC or trastuzumab resistance in HER2+ BC.

A prior study investigating MRV in HER2+ BC cells did not observe a difference in susceptibility to MRV infection between non HER2 expressing and HER2 expressing cells [47]. However, they found that MRV infected and induced cytopathic effects in BC cells that express HER2 (SK-BR-3, KPL4, MDA-MB-453 and CRL1500) and in BC cells that did not express HER2 (MCF7, MDA-MB-231 and Hs578Bst) [47]. To further investigate the impact of MRV infection on HER2+ BC cells, we examined three different cell lines with variable levels of HER2 expression, MCF7 (ER+, PR+/-, HER2 0, Ki-67 90%), ZR-75-1 (ER+, PR+/-, HER2 2+, Ki-67 80%), and AU-565 (ER-, PR-, HER2 3+, Ki-67 90%) [48] for their capacity to support MRV infection and the impact of MRV infection on cell death and relevant cell proliferation pathways. While we found clear evidence that MRV infects and kills BC cells expressing varying levels of HER2, we found little significant change in HER2, Akt, pAkt-Thr308, pAkt-Ser473 or 4E-BP1 during MRV infection that correlated with HER2 overexpression. Because of prior findings that show MRV inhibits the accumulation of HIF-1α, paired with the known relationship between HER2 and HIF-1α, we additionally examined HIF-1α expression following MRV infection in multiple BC cell lines.

## Materials and Methods

### Cells, viruses, antibodies and reagents

AU-565 and ZR-75-1 cell lines were maintained in Roswell Park Memorial Institute (RMPI) 1640 medium (Gibco) supplemented with 10% fetal bovine serum (Atlanta Biologics) and penicillin (100 I.U./ml) streptomycin (100 μg/ml) solution (Mediatech). The MCF7 cell line was maintained in Minimum Essential Media (Gibco) supplemented with 10% fetal bovine serum (Atlanta Biologics), 1% human insulin (Sigma-Aldrich), and penicillin (100 I.U./ml) streptomycin (100 μg/ml) solution (Mediatech). L929 cells were maintained in Joklik modified minimum essential medium (Sigma-Aldrich) supplemented with 2% fetal bovine serum (Atlanta Biologics), 2% bovine calf serum (HyClone), 2 mM L-glutamine (Mediatech), and penicillin (100 I.U./ml) streptomycin (100 μg/ml) solution (Mediatech). Our laboratory stock of MRV strain type 3 Dearing Cashdollar (T3D) was obtained from the laboratory of M. L. Nibert and originated from the lab of L.W. Cashdollar [49]. The virus was propagated and purified as previously described using Vertrel XF (DuPont) instead of Freon [50, 51]. Primary antibodies used were as follows: monoclonal rabbit α-Akt(pan)(11E7) (Cell signaling;#4685), monoclonal rabbit α-Phospho-Akt(Thr308)(D25E6)XP (Cell Signaling; #4056), monoclonal rabbit α-Phospho-Akt(Ser473)(D9E)XP (Cell Signaling;#4060), polyclonal rabbit α-β-actin (Cell Signaling; #4967), monoclonal rabbit α-α-Tubulin (11H10) (Cell Signaling; #2125), polyclonal rabbit α-μNS as described in [52], monoclonal mouse (3E10) α-σNS (University of Iowa, Developmental Studies Hybridoma Bank) [53], monoclonal rabbit α-HER2 (Cell Signaling;# D8F12), monoclonal rabbit α-4E-BP1(53H11) (Cell Signaling;# 9644), monoclonal mouse α-HIF-1α (BD Biosciences; #610958). Secondary antibodies used were Alexa 488- and 594- conjugated donkey α-mouse or α-rabbit IgG antibodies (Invitrogen Life Technologies; #A-21202, #A-21207), and goat α-mouse or α-rabbit IgG AP-conjugate antibodies (Bio-Rad Laboratories, #1706520, #1706518).

### Hypoxia and CoCl_2_ treatment

To mimic hypoxic tumor environments, AU-565, MCF7 and ZR-75-1 were incubated in Galaxy 48R CO_2_ Incubator (New Brunswick Scientific) equipped with 1-19% O_2_ control set at 1% O_2_, 5% CO_2_, and 37°C for 4 hours, or treated with 100mM Cobalt (II) Chloride (CoCl_2_) for 4 hours prior to harvesting.

### Infection

24 hours post-plating (h p.p.), AU-565, MCF7, and ZR-75-1 were infected with T3D in phosphate-buffered saline (PBS) (137 mM NaCl, 3 mM KCl, 8 mM Na_2_HPO_4_, 1.5 mM KH_2_PO_4_, pH 7.4) with 2 mM MgCl_2_ for 1 h at 37°C with shaking every 15 minutes. Media was replenished and infected cells incubated at 37°C with 5% CO_2_. For experiments using serum starvation, media was removed 23 hours post-infection (h p.i.), cells were washed with PBS, then media without FBS was added for the final hour of infection. Multiplicity Of Infection (MOI) for each cell line was determined by first measuring cell infectious units (CIU) on each cell line, as described in [52]. Once CIU/ml of the viral preparation was determined, the amount of virus to add to cells was calculated by multiplying the number of cells by the desired MOI, which gives the total CIU needed to infect at that MOI.

### Immunoblotting

AU-565, MCF7, and ZR-75-1 were plated at 2 x 10^5^ per well in 12 well plates. 24 h p.p., cells were mock-infected or infected with T3D at an MOI of 20 and at 24 h p.i. cells were washed in PBS and collected in 2X protein loading buffer (125 mM Tris-HCL [pH 6.8], 200 mM dithiothreitol [DTT], 4% sodium dodecyl sulfate [SDS], 0.2% bromophenol blue, 20% glycerol). Cell lysates were heated at 95°C for 10 minutes before being separated by sodium dodecyl sulfate-polyacrylamide gel electrophoresis (SDS-PAGE) and then transferred onto nitrocellulose by electroblotting. Nitrocellulose membranes were blocked with Tris-buffered saline [20mM Tris, 137 mM NaCl [pH 7.6]) with 0.1% Tween 20 (TBST)] and 5% milk for 15 minutes, incubated with primary and secondary antibodies in TBST and 5% milk for 18 h (primary) and 4 h (secondary) with 3X PBS washes for 15 minutes after each incubation, followed by addition of NovusBiologicals NovaLume Atto Chemiluminescent Substrate (AP) (Fisher scientific, NBP2-61914). Chemiluminescence images were captured using a ChemiDoc XRS Imaging System (Bio-Rad).

### Immunofluorescence

AU-565, MCF7, and ZR-75-1 were plated on glass coverslips at 2 x 10^5^ in 12-well plates. 24 h p.p., cells were infected with T3D at an MOI of 10. 24 h p.i. cells were fixed with 4% formaldehyde for 20 min, permeabilized with 0.2% Triton X-100 in PBS for 5 minutes and blocked with 1% bovine serum albumin in PBS (PBSA) for 10 minutes with 3 PBS washes between each step. Cells were then incubated for 1 hour at room temperature with primary antibodies diluted in PBSA, washed 3 times with PBS, followed by incubation with secondary antibodies diluted in PBSA for 1 hour before 3 additional PBS washes. Coverslips with labeled cells were mounted with ProLong Gold antifade mountant with DAPI (4,6-diamidino-2-phenylindole dihydrochloride) (Invitrogen, P36931) on glass slides. Each coverslip was then examined on a Zeiss Axiovert 200 inverted microscope equipped with fluorescence optics to either detect viral proteins or determine the number of cells expressing nuclear HIF-1α. To quantify the number of cells expressing HIF-1α in their nucleus, we counted the total amount of cells using DAPI staining, and the number of cells expressing HIF-1α in 10 separate fields of view within each sample. Representative images were captured with a Zeiss AxioCam MR color camera using AxioVision software (4.8.2). Images were prepared using Photoshop and Illustrator software (Adobe Systems).

### Replication assay

AU-565, MCF7, or ZR-75-1 were plated at 1 x 10^6^ per well in 6 well plates. Twenty-four h p.p., cells were infected with T3D at an MOI of 0.1 for 1 hour with gentle rocking every 15 minutes. Media was replenished and cells were subjected to 3 freeze thaw cycles at 0, 6, 12, 18, 24, 30, 36, 42 and 48 h p.i. for AU-565, MCF7, and ZR-75-1. Samples were subjected to serial 1:10 dilutions in PBS with 2mM MgCl_2_. 100μl of samples were added to confluent monolayers of L929 cells, plated at 1.2 x 10^6^ per well in 6-wells 24 hours prior and virus absorption continued for 1 hour at room temp with rocking every 10-15 minutes. After 1 hour, L929 monolayers were overlaid with 2ml of m199 (Irvine Scientific) containing 2 mM L-glutamine (Mediatech), and penicillin (100 I.U./ml) streptomycin (100 μg/ml) solution (Mediatech), 0.225% NaHCO_3_, 2mg/ml trypsin and 1% Bacto-Agar (BD, 214010) and incubated for 3 days at 37°C and 5% CO_2_. Plaques were counted at day 3, made relative to the 0-hour timepoint and graphed using Excel (Microsoft).

### Apoptosis detection

AU-565, MCF7, or ZR-75-1 were plated at 1 x 10^4^ per well in 96 well plates. Twenty-four h p.p., cells were mock-infected or infected with T3D at an MOI of 1. 0, 24, 48, or 72 h p.i. cells were incubated with ApoTox-Glo^TM^ Triplex Assay (Promega, #G6320) according to manufactures recommendation. Fluorescence was quantified using a Glomax Multi Detection Plate Reader (Promega).

### Statistical analysis

For replication assay graphs, error bars were generated by calculating standard deviation using =stdev() function in Excel. All other graphs show mean, as larger line, and one standard deviation on either side of mean as the smaller lines, both calculated using JMP. P-values were calculated using a 2 sided student’s T test, assuming equal variance using =ttest() function in Excel.

## Results

### MRV replication in BC cells in vitro is not influenced by HER2 expression levels

MRV has been shown to infect and replicate in a variety of tumor cell types, including BC cells expressing, or not expressing HER2 [47], however little is known about how HER2 overexpression might influence MRV growth. Three cell lines with varying HER2 protein expression levels were used in our study to determine the impact of HER2 overexpression on MRV infection. MCF7 cells exhibit the lowest HER2 expression, ZR-75-1 cells possess intermediate HER2 expression, and AU-565 cells overexpress HER2 (Figs 1A and 3). To initially confirm that each cell line was susceptible to MRV infection, cells were infected at an MOI of 10 with the T3D serotype of MRV, which is identical in amino acid (aa) sequence to the virus currently used in clinical trials [54], and subjected to immunofluorescence assay using antibodies against the viral non-structural protein, σNS, that is only expressed during active replication. As seen in Fig 1A, σNS is expressed in each of the tested cell lines suggesting that HER2 expression does not inhibit MRV entry, transcription, or translation. To quantify the extent by which MRV replication may be influenced by cellular HER2 expression levels, we performed replication assays. Each cell line was infected with T3D at an MOI of 0.1 and samples were collected every 6 hours for a total of 48 hours. Following freeze/thaw processing, samples were analyzed via plaque assay on L929 cells to determine viral titer at each time point. As seen in Fig1B, the T3D replication curve is similar irrespective of HER2 status and after 48 hours each cell line showed an increase in viral titer of three logs, which suggests that MRV replication is not affected by HER2 expression levels.

**Fig 1.**
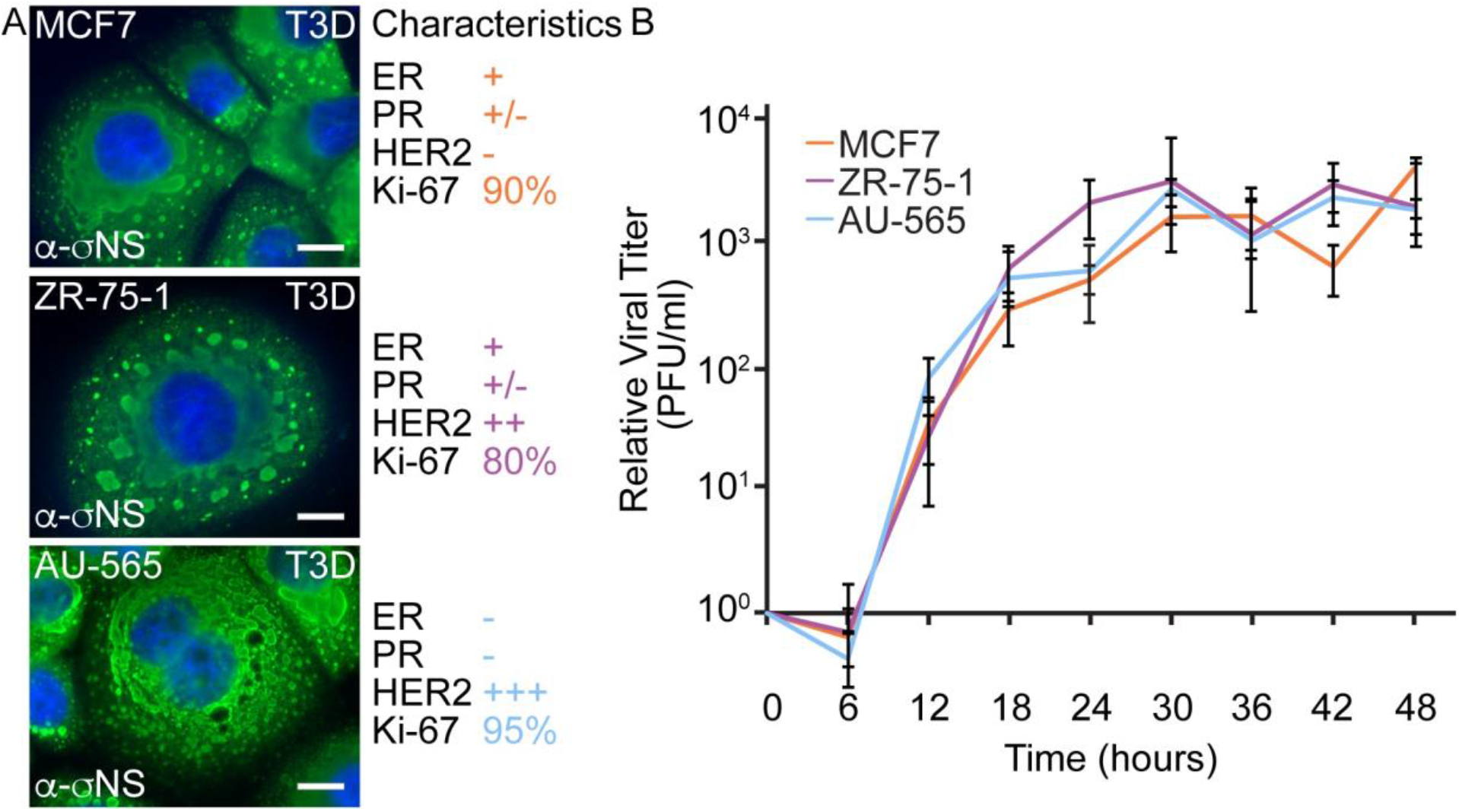
MRV replicates efficiently in BC cells expressing variable levels of HER2. A) MCF7 (top), ZR-75-1 (middle) and AU-565 (bottom) cells were infected with T3D and at 24 hours p.i. were fixed and immunostained with mouse α-σNS monoclonal, followed by Alexa 488-conjugated donkey α-mouse IgG (green), and counter stained with DAPI (blue). Scale Bars=10 μm. The ER, PR, HER2, and Ki-67 status of each cell line is shown. B) MCF7, ZR-75-1 and AU-565 cells were infected with an MOI of 0.1 and at 0, 6, 12, 18, 24, 30, 36, 42, and 48 h p.i. samples were analyzed for viral titer at each timepoint via plaque assays on L929 cells. The means and standard deviation from three experimental replicates are shown.

### MRV induces BC cell cytotoxicity in part through apoptosis

MRV has been shown to induce cell death via apoptosis and necroptosis in various types of cancer cells [32, 47, 55-57]. To determine the extent that MRV infection induces cell death in MCF7, ZR-75-1 and AU-565, cells were infected with T3D at an MOI of 1 or mock infected and analyzed simultaneously for cell viability and cytotoxicity at 0, 24, 48, and 72 h p.i. using the Apo-Tox Glo assay which measures the cleavage products of cell-permeant GF-AFC substrate, indicating live cells, and cell-impermeant bis-AAF-R110 substrate, indicating dead cells. These experiments show a significant decrease in cell viability in all infected samples as compared to mock starting at 24 h p.i. in MCF7, 72 h p.i. in ZR-75-1 and 48 h p.i. in AU-565 (Fig 2A). Additionally, we saw a significant increase in cytotoxicity starting at 24 h p.i. for MCF7 and AU-565 (Fig 2B). The ZR-75-1 cell line did not show significant cytotoxicity at any of the timepoints, however that is not unexpected since the decrease in viability for these cells was delayed and did not occur until 72 h p.i.

**Fig 2.**
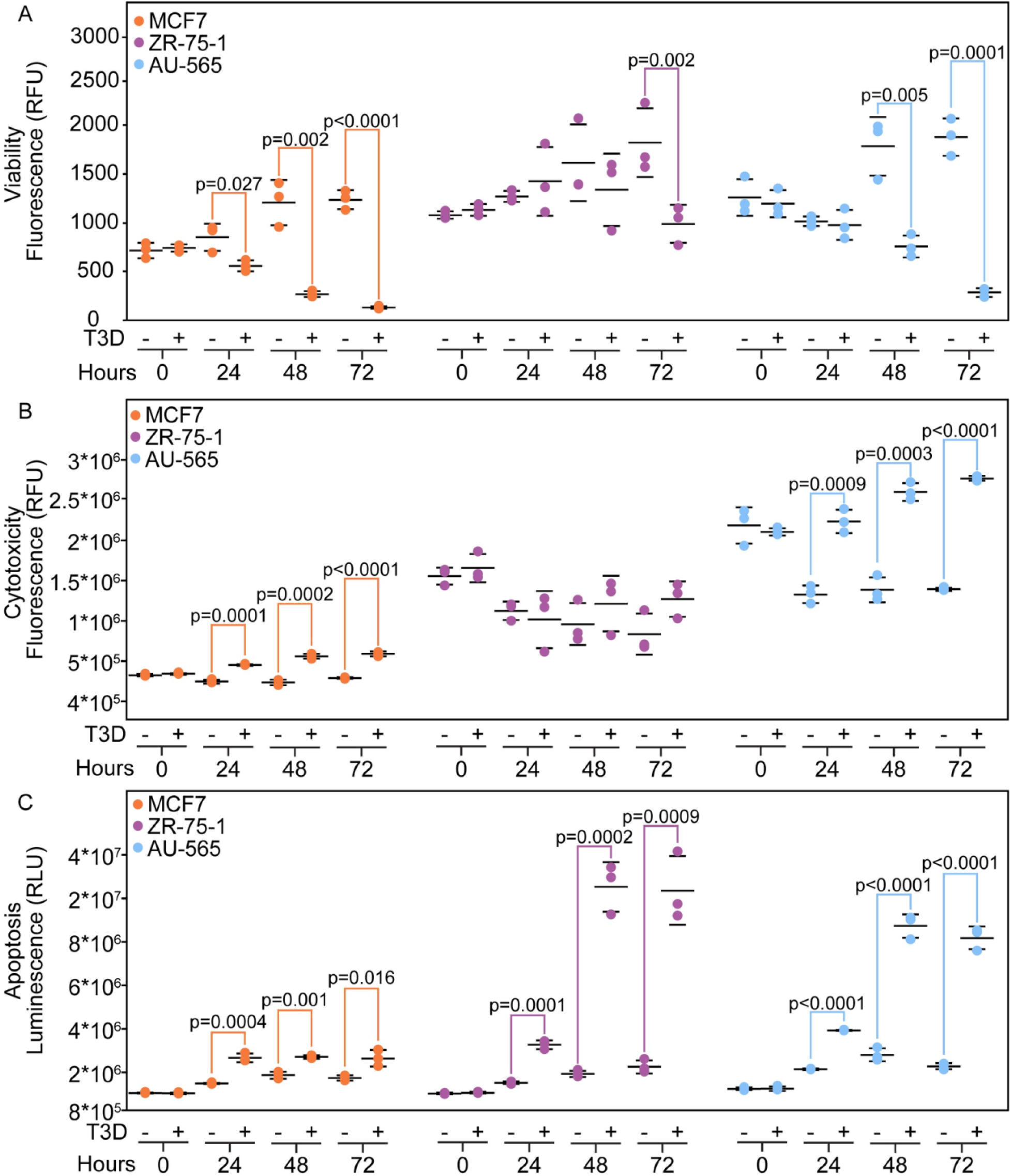
MRV infection induces BC cell death via caspase 3/7 dependent apoptosis. MCF7, ZR-75-1, and AU-565 cells were mock-infected or infected with T3D at an MOI of 1 and at 0, 24, 48, or 72 h p.i. samples were subjected to A) Viability assay, B) Cytotoxicity assay, and C) Caspase 3/7 activation assay. Means and standard deviation of 3 experimental replicates are shown, and significant differences between samples are indicated. P-values were calculated using student’s 2-sided T-test and are shown.

To begin to examine the mechanism of MRV-induced cell death, we measured caspase 3/7-dependent apoptosis in the BC cell lines following MRV infection using a proluminescent caspase-3/7 DEVD-aminoluciferin substrate and a thermostable luciferase (Caspase-Glo®3/7). MCF7, ZR-75-1, and AU-565 cells were infected with T3D at an MOI of 1 or mock-infected and at 0, 24, 48 or 72 h p.i., caspase activation was measured. Despite the variation in times p.i. that MRV caused a reduction in cell viability or increase in cytotoxicity, all three cell lines showed significant increase in caspase 3/7 activity in infected cells compared to mock starting at 24 h p.i. which increased relative to mock-infected cells over time (Fig 2C). Taken together with the cytotoxicity/viability data, this shows that MRV infection induces cell death in part via induction of apoptosis in BC cells, and that overexpression of HER2 does not appear to influence apoptosis induction or cell death by MRV.

### MRV infection does not significantly alter the PI3K/Akt pathway in HER2 overexpressing BC cells

One of the primary downstream pathways that is activated by HER2 is the PI3K/Akt pathway. Activation of this pathway allows cancer cells to override apoptosis signals, increases cell growth, and increases tumor cell proliferation [6]. As MRV infected cells have previously been shown to have reduced pAkt-Ser473 [33, 37, 58], depending on MRV serotype and cell type, we investigated the capacity of infection to modify HER2 expression and the HER2-activated downstream PI3K/Akt pathway. MCF7, ZR-75-1, and AU-565 cells were mock-infected or infected with T3D at an MOI of 20. At 24 h p.i. samples were collected, and cell lysates were separated on SDS-PAGE and transferred to nitrocellulose, then immunoblotted using cell-protein specific antibodies. Upon examination of HER2 expression in uninfected and infected cells we found that while there was a consistent decrease in HER2 levels in MRV infected cells, over repeated experiments the expression of HER2 protein was not significantly influenced by MRV infection (Fig 3).

**Fig 3.**
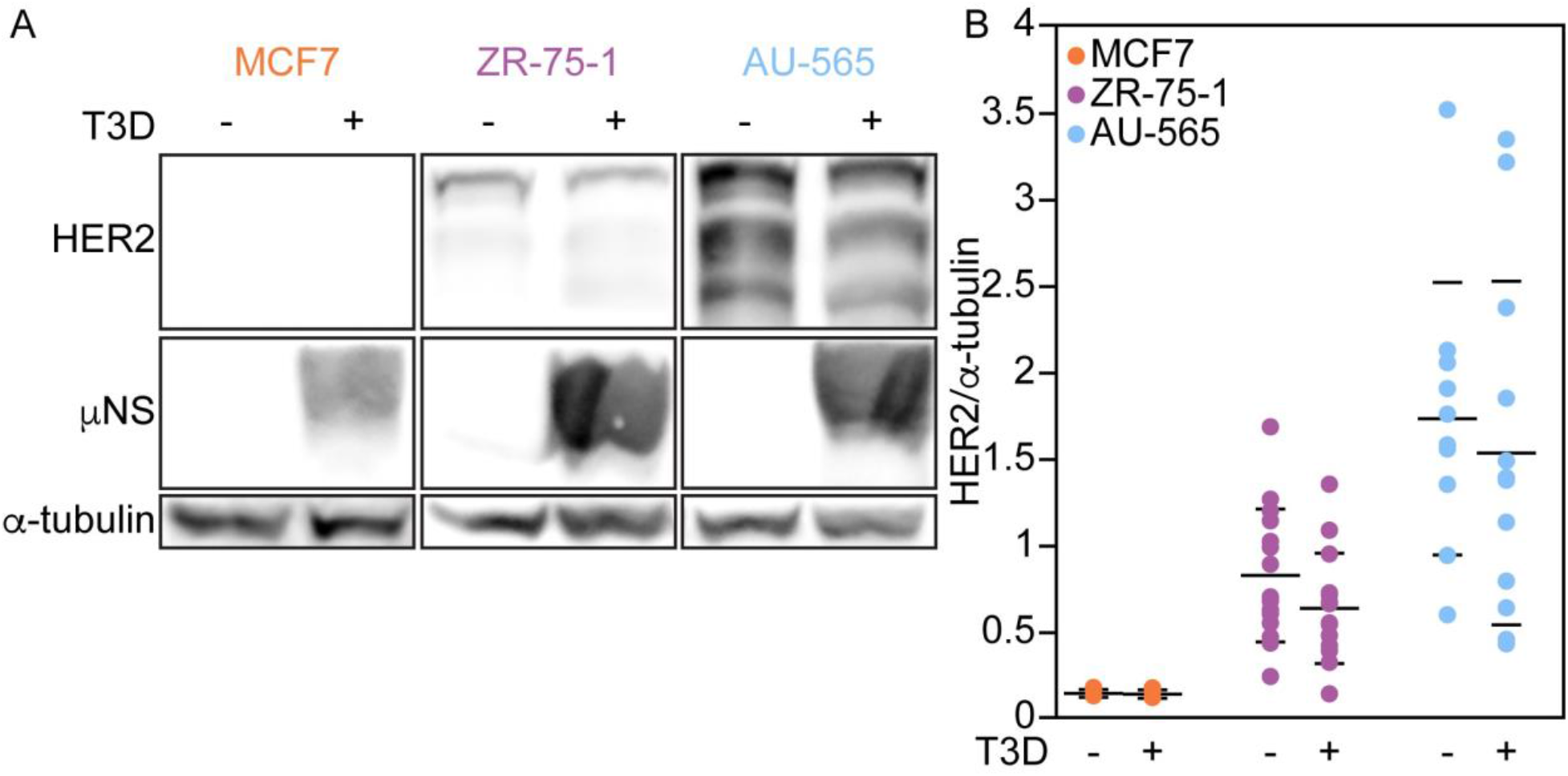
HER2 expression is not significantly altered during MRV infection. A) MCF7, ZR-75-1, and AU-565 cells were mock-infected or infected with T3D at an MOI of 20. At 24 h p.i. samples were collected and subjected to immunoblot using monoclonal rabbit α-HER2, polyclonal rabbit α-μNS and polyclonal rabbit α-β-actin antibodies followed by goat α-rabbit IgG-AP conjugate secondary antibody. Proteins were detected using AP chemiluminescence substrate. B) Immunoblots were analyzed using Quantity One software to quantify HER2 expression levels relative to β-actin. Means and standard deviation from at least three biological replicates with three experimental replicates are shown. P-values were calculated using student’s 2-sided T-test and are shown.

Despite non-significant decreases in HER2 expression following MRV infection, the possibility remained that proteins downstream of HER2 in the PI3K/Akt pathway may be altered by MRV, therefore we also investigated expression levels of total Akt, 4E-BP1, and of phosphorylated Akt at two known regulatory sites. We found a significant decrease in total Akt expression in MCF7 cells, but not in the HER2 cell lines (Fig 4A, B). Moreover, both the Ser(473) and Thr(308) phosphorylation sites on Akt remained unchanged during MRV infection in the HER2 expressing cell lines, while MCF7 cells demonstrated a decrease in pSer473Akt during MRV infection (Fig 4C-F). 4E-BP1, one of the downstream effector proteins in the PI3K/Akt pathway, remained unchanged during infection of MCF7, ZR-75-1 and AU-565 (Fig 4G, H). Despite the decrease in Akt and pAkt-Ser473 in MCF7 cells during MRV infection, 4E-BP1 protein levels in mock and infected cells were also the same in these samples, suggesting MRV infection does not alter this downstream target of the PI3K/Akt pathway in these cells. It is possible that the decrease in Akt and pAkt-Ser473 results in modulation of other proteins, such as mouse double minute 2 homolog (MDM2), p27, Bcl-2 antagonist of cell death (BAD), IkB kinase (IKK), glucose transporter type (GLUT) or others since activated Akt is pivotal in multiple regulatory pathways [59]. Taken together, these results show that T3D infection does not substantially modulate the HER2/PI3K/Akt pathway in cells expressing or overexpressing HER2.

**Fig 4.**
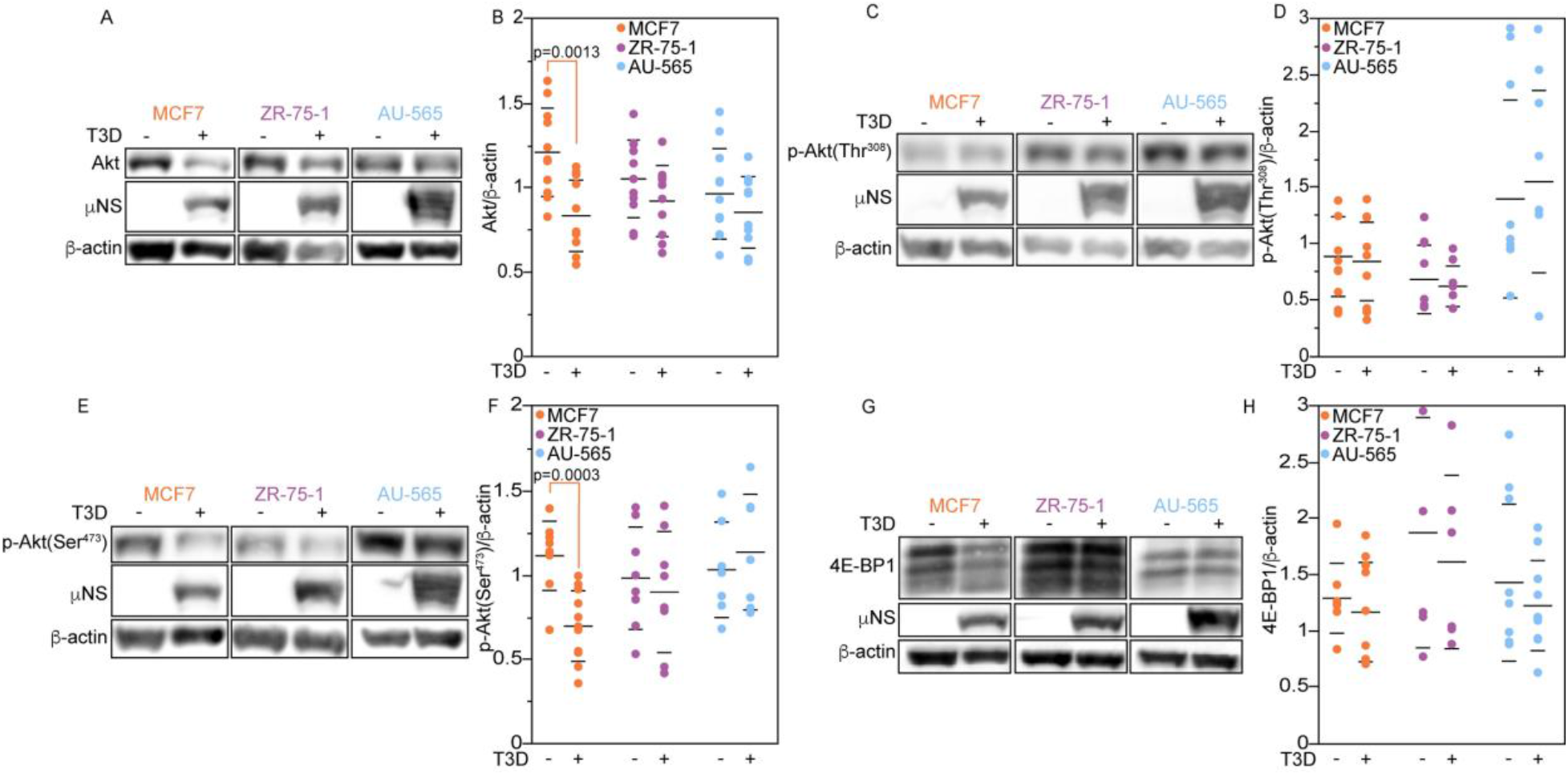
MRV does not alter the PI3K pathway in HER2 expressing cells. MCF7, ZR-75-1, and AU-565 cells were mock-infected or infected at an MOI of 20 with T3D. At 23 h p.i. media was removed, and serum free media was added for 1 hour. Cells were collected and subjected to immunoblot analysis using polyclonal rabbit α-μNS antibodies, polyclonal rabbit α-β-actin antibodies, and A/B) monoclonal rabbit α-Akt(pan) antibody, C/D) monoclonal rabbit α-phospho-Akt(Thr308) antibody, E/F) monoclonal rabbit α-phospho-Akt(Ser473) antibody, or G/H) monoclonal rabbit α-4E-BP1 antibody, followed by goat α-rabbit IgG-AP conjugate secondary antibody. Chemiluminescence from immunoblots was analyzed using Quantity One software and means and standard deviation from multiple biological replicates are shown. P-values were calculated using student’s 2-sided T-test in Excel and are shown in samples with significant differences.

### MRV inhibits accumulation of HIF-1α under hypoxic conditions in BC cells

In prostate, lung, colon, and sarcoma tumor models MRV has been shown to decrease the accumulation of HIF-1α in hypoxic conditions [30-32, 35, 37]. Interestingly, HIF-1α expression in implicated in progression to trastuzumab resistance. Moreover, HER2 overexpression has been shown to increase accumulation of HIF-1α in normoxic conditions [44, 45]. We hypothesized that HER2 overexpression may impact the effect of MRV on HIF-1α. To examine HIF-1α levels in HER2 negative and positive BC cells, MCF7, ZR-75-1, and AU-565 cells were mock-infected or infected with MRV at an MOI of 20. Infection was allowed to continue for 20 hours, at which point half of the samples were incubated in 1% oxygen for the final 4 hours of infection, while the other half remained in normoxic conditions. Samples were collected and cell lysates were subjected to separation via SDS-PAGE, transferred to nitrocellulose, and immunoblotted using HIF-1α antibody. In these experiments, MRV infection significantly decreased the amount of HIF-1α accumulation in cells grown under hypoxic conditions in all BC cell types (Fig 5A, B). Unlike previous reports [44, 45], we did not measure increased accumulation of HIF-1α in HER2 expressing cell lines grown under normoxic conditions (Fig 5A). To confirm these results and examine HIF-1α expression and localization to the nucleus at the individual cell level, BC cells were mock-infected or infected with MRV at an MOI of 20. At 20 h p.i. 100 mM CoCl_2_, which mimics hypoxia by inhibiting PHD2 hydroxylation of HIF-1α [60] and stabilizes HIF-1α under normoxic conditions, was added for 4 h, at which point samples were fixed, permeablized, and immunostained with antibodies that bind HIF-1α and viral non-structural protein μNS. Under CoCl_2_ treatment, in mock-infected cells, HIF-1α expression is increased relative to untreated cells, and the protein is localized to the nucleus. However, in MRV infected cells there is significantly less HIF-1α accumulation or nuclear localization (Fig 5C, D). Taken together, the immunoblot and immunofluorescence data provide clear evidence that under hypoxic conditions MRV reduces accumulation of HIF-1α in BC cells and that HER2 overexpression does not alter this phenotype.

**Fig 5.**
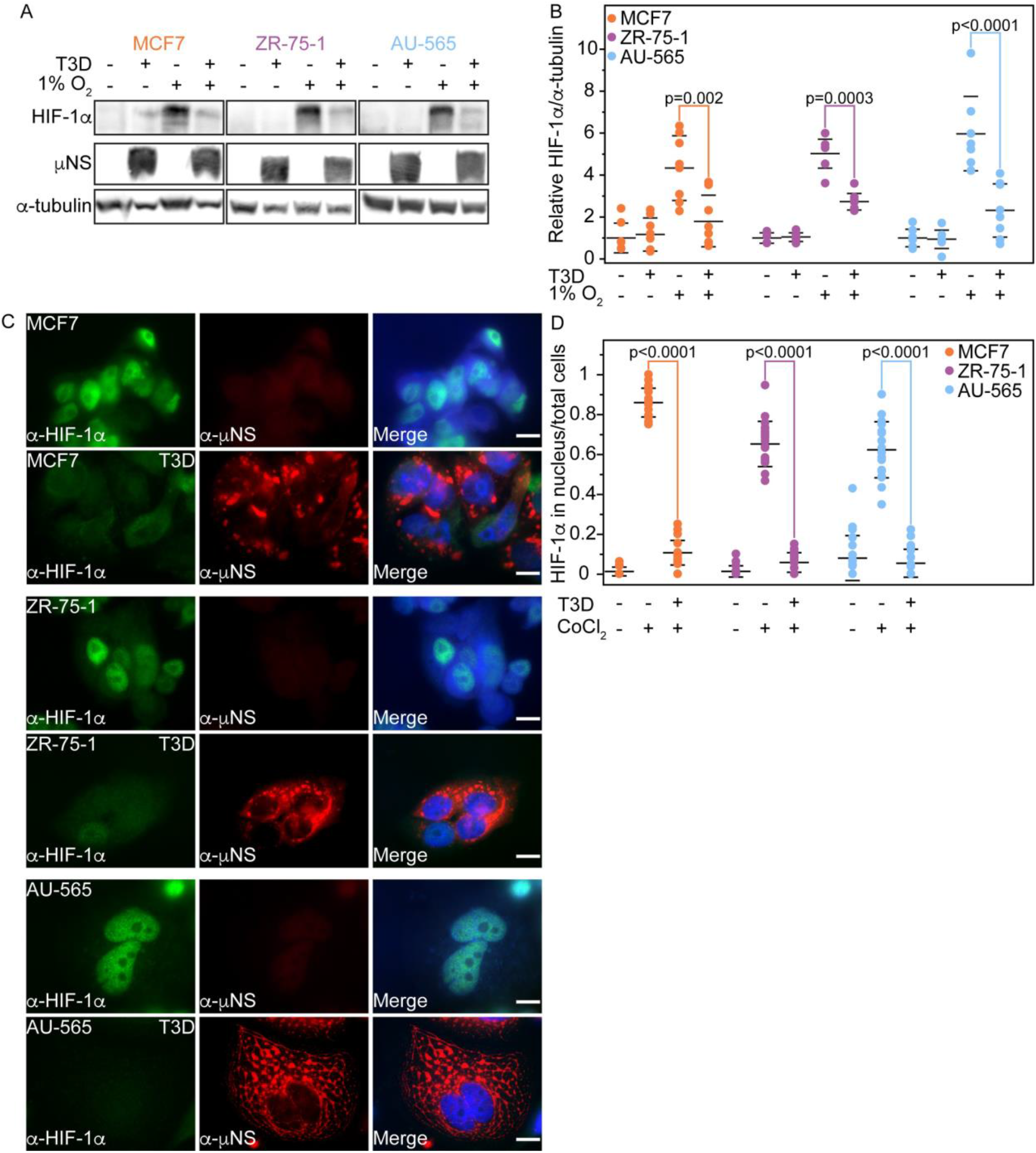
MRV infection inhibits accumulation of HIF-1α in BC cells. MCF7, ZR-75-1, and AU-565 were mock-infected or infected with T3D at an MOI of 20. A) At 20 h p.i. cells were subjected to normoxic (20% oxygen) or hypoxic conditions (1% oxygen) for 4 hours then samples were collected and subjected to immunoblot analysis using monoclonal mouse α-HIF-1α antibody, polyclonal rabbit α-μNS antibodies, and monoclonal rabbit α-α-tubulin antibody followed by goat α-rabbit IgG AP-conjugate secondary antibody or goat α-mouse IgG AP-conjugate secondary antibody and chemiluminescence imaging. B) Quantification of chemiluminescence from multiple biological replicates from A. Mean and standard deviation are shown and P-values are indicated. C) At 20 h p.i., cells were subjected to 100 mM CoCl_2_ for 4 hours then fixed and immunostained with monoclonal mouse α-HIF-1α antibody (left) and polyclonal rabbit α-μNS antibodies (right) followed by Alexa 488-conjugated donkey α-mouse (green) and Alexa 594-conjugated donkey α-rabbit secondary antibodies (red). Cells were counterstained with DAPI (blue) and merged images are shown. Bar=10μM D) 10 fields of view from C were analyzed per sample for each biological replicate to quantify HIF-1α expression. Mean and standard deviation are shown, and P-values calculated using student’s 2-sided T test are indicated.

## Discussion

This study confirms that MRV infects and replicates in various subtypes of BC cells and shows that HER2 overexpression does not interfere with or enhance the ability of MRV to replicate. Despite robust replication, we did not observe a significant impact from MRV infection on multiple proteins in the PI3K/Akt downstream pathway in our experiments that correlated with HER2 expression levels. It is possible that MRV may transiently alter these proteins, and that our experiments, which were performed at 24 h p.i., did not capture these changes. We did measure a robust decrease in HIF-1α accumulation in MRV infected BC cells independent of HER2 expression levels, which provides further evidence that MRV alters this important hypoxic response pathway.

Though most of the results examining MRV-induced tumor cell viability, cytotoxicity, and apoptosis were congruent, in the ZR-75-1 cell line, which expresses moderate levels of HER2, caspase 3/7 activity was evident at every time point starting at 24 h p.i., but viability was not significantly decreased until 72 h p.i. and there was no significant difference in cytotoxicity at any time point. These differences can likely be explained by the limitations of the cytotoxicity assay. The substrate added to detect cytotoxicity requires that cell membranes are degraded to allow active dead-cell protease into the supernatant for cleavage to occur and fluorescence to be detected. It is possible for caspase 3/7 activity to be present without detectable cytotoxicity [61]. In AU-565 cells, which express very high levels of HER2, there was an increase in cytotoxicity at 24 h p.i., suggesting that membrane degradation may occur more rapidly in AU-565 infected cells compared to ZR-75-1 infected cells.

In previous studies MRV has been shown to modulate the amount of pAkt-Ser473, depending on the cell line tested [32, 33, 37, 58]. The prostate cancer cell line, LNCaP [33], the stomach cancer cell line, SNU-216 [37], and three canine mast cell tumor cell lines, VIMC, CoMS, and CMMC [58] have all shown decreased pAkt-Ser473 during MRV infection, however one canine mast tumor cell line, HRMC showed no change in pAkt-Ser473 during MRV infection [58]. Here we show that in ZR-75-1 and AU-565 cells MRV infection does not change the levels of pAkt-Ser473, pAkt-Thr308, or total Akt. However, in MCF7 cells, MRV infection results in decreased pAkt-Ser473 and total Akt. This work provides additional evidence that the decrease in pAkt-Ser473 due to MRV infection is cell line dependent. Alternatively, it is possible that expression of HER2 in ZR-75-1 and AU-565 leads to increases in total Akt expression and pAkt-Ser473 phosphorylation that override MRV infection-induced decreases. Nonetheless, the decrease in pAkt-Ser473 did not result in significant changes in 4E-BP1 expression, a downstream protein in the mTOR pathway. This is not surprising as phosphorylation at the Thr308 of Akt is necessary for 4E-BP1 to be downregulated and pAkt-Thr308 remained unaffected during MRV infection [62]. Overall, our results suggest that MRV infection does not alter downstream signaling of the HER2 pathway in cells overexpressing HER2 but may have an impact on Akt phosphorylation and accumulation in the absence of HER2 in MCF7 cells.

Approximately 24-56% of invasive BC cells exhibit hypoxic regions [38, 40, 41, 63]. In these areas, accumulation of HIF-1α and activation of downstream target genes of the hypoxic response causes increased growth and brain metastasis [64]. Our lab and others have shown HIF-1α accumulation is inhibited in hypoxic environments in various cell lines, and *in vivo* during MRV infection [29-32, 34]. This study supports previous work and adds BC cells to the growing number of cancer types in which HIF-1α is prevented from accumulating in hypoxic conditions during MRV infection. There are several cancer therapeutics, including radiotherapy, some chemotherapies, and immunotherapies that are inhibited to varying degrees in hypoxic tumor microenvironments [65-67]. In addition, novel therapeutics are being developed specifically to inhibit HIF-1α [68-73]. Finally, recent data provides evidence that HIF-1α expression is responsible for progression to resistance to trastuzumab in a HER2+ BC cell line, SKBR3 [15]. Signal transducer and activator of transcription 3 (STAT3) regulates both transcription and nuclear localization of HIF-1α, leading to elevated transcription factor, hairy and enhancer of split 1 (HES-1) expression which in turn lowers levels of phosphatase and tensin homolog (PTEN) in trastuzumab resistant SKBR3 [15]. Additionally, trastuzumab resistant SKBR3 showed a response to trastuzumab when siRNA was used to knock down HIF-1α, suggesting expression of HIF-1α is a major contributor to trastuzumab resistance [15]. The data from this study provides further evidence that MRV therapy may enhance the efficacy of BC drugs/therapies that are inhibited by or target hypoxia.

MRV has been shown to have the most clinical benefit when used in combination with a second therapeutic [19], therefore it would be worth testing the effect of dual MRV and HER2+ BC specific treatments to determine if MRV works synergistically with other therapies to kill HER2+ BC cells. In HER2 overexpressing gastric cancer cells and a mouse model, MRV enhanced trastuzumab’s antitumor efficacy, suggesting MRV has the potential to work synergistically in combination with this HER2-specific therapeutic [74]. A clinical trial investigating MRV use with trastuzumab in patients with HER2+ BC is currently in progress [75]. Increasing our understanding of how MRV infection influences other HER2+ BC-specific therapeutics in vitro and in vivo will inform ongoing and future clinical trials.

Overall, this study provides additional pre-clinical evidence that MRV has potential as a therapeutic option against HER2+ BC. While the virus does not appear to inhibit HER2-activated pathways, we have added to prior evidence in different cell lines that HER2 overexpression does not inhibit MRV replication and tumor cell death [47, 76]. Though the mode of action of MRV has historically been characterized as tumor cell lysis, it is increasingly clear that activation of the patient immune system against MRV infected tumor cells is of equal, if not more, importance. In this regard, our data showing MRV-induced tumor cell death in BC cells overexpressing HER2 would suggest that in addition to the recruitment of inflammatory cytokines, and induction of an anti-viral immune response against infected tumor cells, MRV killing of these cells would induce the robust release of HER2 antigens, which would then activate an anti-HER2 tumor response. Although we were focused on the influence of HER2 overexpression, our data also indicates MRV readily infects and kills BC cells with ER+, PR+/- genetic signatures.

The importance of molecular genetic subtyping and differences in tumor microenvironment between patients diagnosed with major tumor cell types is being recognized as increasingly important in the search for effective and curative therapeutics. Modalities designed to target specific genetic subtypes of BC, particularly HER2+ BC, have added valuable long-term survival time to patients receiving them, supporting the importance of moving beyond cell type specific therapies into a more personal molecular genetic approach. Combinatorial trials including MRV with specific therapeutics against known BC genetic subtypes are underway. For example, a clinical trial (NCT04102618) investigating MRV use in various subtypes of BC as a dual therapy administered currently available FDA approved BC therapies such as letrozole, atezolizumab, and trastuzumab, dependent on BC subtypes, with MRV (Biotech and SOLTI, 2022). Results from this trial are still being collected and have not yet been published but will provide insight into how targeting specific genetic BC subtypes (eg. atezolizumab against PD-L1, trastuzumab against HER2) perform when administered in combination with MRV. Furthering our understanding of MRV infection of specific molecular genetic tumor subtypes in pre-clinical models such as that presented in this study will inform more targeted treatment, determine if the therapies are most effective when given together or in sequence, and provide important information that may allow insight into future clinical trial design.

## Acknowledgments

We would like to acknowledge the fellow members of the Miller laboratory for their input and discussion. This work was supported by the Margaret B. Barry Research Award, the NIH R15CA202984 grant, and Lloyd Cancer Research Fund.

